# Hexa-acylated lipopolysaccharides from the gut microbiota enhance cancer immunotherapy responses

**DOI:** 10.1101/2024.06.24.600291

**Authors:** Benjamin S. Beresford-Jones, Puspendu Sardar, Wangmingyu Xia, Omar Shabana, Satoshi Suyama, Ruben J. Jesus Faustino Ramos, Amelia T. Soderholm, Panagiotis Tourlomousis, Paula Kuo, Alexander C. Evans, Charlotte J. Imianowski, Alberto G. Conti, Alexander J. Wesolowski, Klaus Okkenhaug, Sarah K. Whiteside, Rahul Roychoudhuri, Clare E. Bryant, Justin R. Cross, Virginia A. Pedicord

## Abstract

Immune checkpoint inhibitors (ICI), such as anti-PD-1, have revolutionized cancer treatment, but they are only effective for a minority of patients. The gut microbiome plays a crucial role in modulating immunotherapy treatment responses, and previous studies correlated lipopolysaccharide (LPS)-producing gut microbes with poorer prognosis. However, LPS from diverse bacterial species have activities ranging from immunostimulatory to inhibitory. By functionally analyzing fecal metagenomes from 112 melanoma patients prior to anti-PD-1 therapy, we found that a subset of LPS-producing bacteria encoding immunostimulatory hexa-acylated LPS was enriched in the microbiomes of clinical responders. We confirmed robust activation of the NF-kB pathway by hexa-acylated LPS *in vitro*, and this activation was significantly inhibited by penta-acylated LPS in a dose-dependent manner. Importantly, oral administration of hexa-acylated LPS augmented anti-PD-1-mediated anti-tumor immunity in an *in vivo* mouse model of cancer immunotherapy. Microbiome hexa-acylated LPS may therefore represent an accessible predictor and potential enhancer of clinical anti-PD-1 immunotherapy responses.

**Statement of significance:** Functional rather than taxonomic profiling of patient gut microbiomes reveals hexa-acylated LPS as a novel biomarker of responsiveness and a targetable pathway for enhancing responses to anti-PD-1, informing future studies and current patient treatment.

## Introduction

Immune checkpoint inhibitors (ICI) unleash the power of a patient’s own immune system to combat cancer. Multiple landmark studies have identified a role for the gut microbiome in modifying ICI treatment response^1,2^, however, there is little consensus between these studies as to which members of the gut microbiota are important for modifying treatment outcomes. More recent studies instead conclude that the relationship between the gut microbiome and ICI clinical response extends beyond the level of species composition^3^, pointing to associations between unfavorable immunotherapy outcomes and lipopolysaccharide (LPS)-producing bacteria in the gut microbiota^4^. However, beyond association, the effects of microbiota-derived LPS remain incompletely defined, and few studies consider the different structures of LPS produced by gut microbes.

Primarily characterized in pathogenic infections for its role in stimulating immune responses to Gram-negative bacteria, LPS activation of host toll-like receptor 4 (TLR4) has complex context-dependent roles in physiology. In the gut, LPS-TLR4 signaling has been shown to both dampen intestinal immune responses to the resident microbiota during states of health ^5^ and modulate systemic autoimmune activity^6^. However, the immunostimulatory effect of LPS depends on its structure, which differs profoundly between bacterial species. Indeed, not all Gram-negative bacteria in the gut harbor all genes within the LPS biosynthetic pathway, resulting in production of a heterogenous pool of LPS structures by different gut microbes. The lipid A region of LPS that is recognized by TLR4, is composed of two glucosamine sugars linked to a variable number of acyl chains. In humans, hexa-acylated LPS potently activates TLR4 and host immunity, whereas hypo-acylated penta- and tetra-acylated LPS poorly activate TLR4 and can antagonize immune activation by hexa-acylated LPS^6–9^. Recent studies have associated some LPS-encoding Gram-negative taxa, such as members of the *Bacteroidaceae* family, with non-response to ICI treatment^4,10,11^. However, taxonomic associations with treatment response may not capture the diversity of LPS structures present and the resulting complexity of functional interactions occurring between gut-derived LPS species and host immune responses.

To determine the potential functional rather than taxonomic basis for Gram-negative bacteria-mediated modification of ICI response, we analyzed LPS biosynthesis genes in fecal metagenomes of cancer patients undergoing ICI with anti-PD-1 therapy. Employing a multi-cohort meta-analysis combining taxonomic and functional annotation, we identify hexa-acylated LPS-encoding gut bacteria as a potential determinant of clinical response to anti-PD- 1 therapy. Using *in vitro* human monocyte/macrophage reporter assays, we demonstrate a TLR4-dependent role for increased ratios of hexa-acylated LPS in activation of the NF-kB pathway. These findings further our understanding of the complex host-microbiome interactions that define response to ICI treatment and advise against the use of inhibitors of LPS-induced TLR4 signaling suggested by some previous studies^12,13^.

## Results

### Patient gut microbiome LPS acylation segregates responders to anti-PD-1 therapy

To explore the functional role of microbiota-derived LPS in responses to anti-PD-1 immunotherapy, we first analyzed baseline gut microbiota composition prior to treatment using metagenomic sequences from a large-cohort study of anti-PD-1-treated melanoma patients^1^. Non-metric multidimensional scaling (NMDS) ordination based on abundance of Gram-negative bacterial genera did not significantly segregate responders from non-responders **(Fig. 1a, left)**. However, NMDS ordination based on function, specifically genes involved in LPS biosynthesis, significantly segregated responders from non-responders and revealed differential abundance of *lpxM*, which mediates lipid A hexa-acylation (**Fig. 1a, right**). Among genes involved in bacterial LPS assembly, *lpxA, B, C, D, H, K*, and *wdtA/waaA* encode genes required for basic LPS assembly and tetra-acylation of lipid A, whereas *lpxL* facilitates LPS penta-acylation, and *lpxJ* and *lpxM* hexa-acylate LPS with differing acyl chain lengths and positions^14,15^.

**Figure 1.**
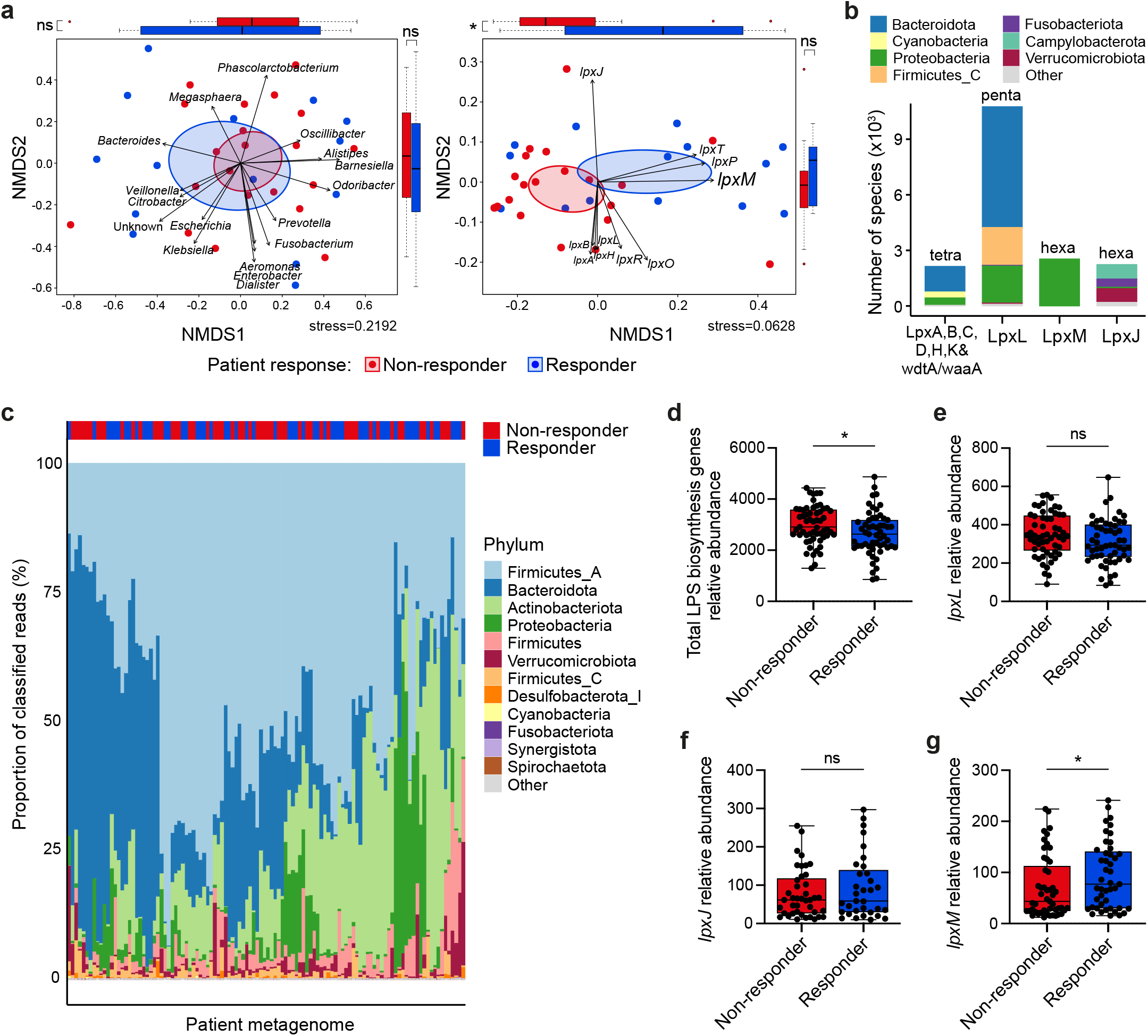
Patient responsiveness to anti-PD-1 therapy is associated with microbiome immunostimulatory hexa-acylated LPS. **a,** Non-metric multidimensional scaling (NMDS) ordination of patient metagenomes with arrows indicating projected bacterial genera (left) and *lpx* encoding genes (right) from melanoma patients prior to anti-PD-1 therapy. n = 35. NMDS stress scores were calculated after 1000 iterations or the best value achieved. Ellipses were drawn with a 95% confidence interval. Boxplots show the distribution of data points (coordinates) within NMDS1 and NMDS2. **b,** Taxonomic breakdown of *lpx* encoding bacteria stacked according to the encoded enzymes for the lipid A tetra-acylated backbone (left) and each level of lipid A acylation as indicated above each stacked bar. **c,** Phylum-level taxonomic composition of pre-treatment fecal metagenomes from 112 melanoma patients. The relative abundance of each phylum is given as a proportion of total classified reads and each phylum is indicated by the fill color of the stacked bars. **d,** Relative abundance as count per million (CPM) of total LPS biosynthesis genes and **e,** *lpxL* (penta-acylation), **f,** *lpxJ* (hexa-acylation) and **g,** *lpxM* (hexa-acylation) genes within fecal metagenomes of responder and non-responder metastatic melanoma patients prior to treatment with anti-PD-1. Mann-Whitney test with bars and error representing median and range (d-g). \**P* < 0.05.

We next expanded our analyses by performing taxonomic and functional annotation on metagenomes from five published datasets encompassing 112 melanoma patients treated with anti-PD-1 from seven clinical studies across four different countries, batch correcting for variance introduced by study and geographic differences **(Extended Data** Fig. 1)^1,3,4,11,16^. Importantly, only samples taken before the start of immunotherapy from patients not taking probiotics, antibiotics or medications like proton pump inhibitors and H2 receptor antagonists were included, as these treatments are known to have significant effects on the microbiome, complicating assessment of the patient’s baseline microbiota^3^. On a taxonomic level, a majority of the commensal Gram-negative species in patient gut microbiotas encoded *lpxL*-mediated capacity for penta-acylated LPS biosynthesis, dominated by members of the Bacteroidota phylum **(Fig. 1b)**, which has previously been demonstrated to be immunoinhibitory^7,8^. While we observed considerable inter-patient variation on a taxonomic level **(Fig. 1c)**, the abundance of total LPS biosynthesis genes was significantly higher in non-responders to anti-PD-1 therapy **(Fig. 1d)**, as previously noted^4^. Amongst the genes involved in modification of lipid A, *pagL* (lipid A 3-O-deacylation), *pagP* (palmitoylation), *lpxE* and *lpxF* (dephosphorylation), *lpxL* (penta-acylation) and *lpxJ* and *lpxM* (different forms of hexa-acylation), only *lpxM* was significantly different between responders and non-responders **(Fig. 1e-g)**, predominantly represented by Proteobacteria (homonym Pseudomonadota) **(Fig. 1b)**. Notably, these functional comparisons of microbiome LPS biosynthesis genes were consistent with those from pooled and per-study data prior to batch correction **(Extended Data** Fig. 2). These analyses confirmed a negative correlation between total gut microbiome LPS and immunotherapy response but uncovered a significant positive association between *lpxM*-mediated LPS hexa-acylation and favorable anti-PD-1 therapy outcomes.

### Functional metagenomic analyses reveal elevated hexa-acylated LPS signatures in immunotherapy responders

To validate our findings using additional unbiased clustering methods, we identified three functional enterotypes (clusters of bacterial communities in the gut) using partitioning around medoids (PAM) clustering with a Jensen-Shannon divergence (JSD) distance metric calculated from the functional annotation profiles of the 112 melanoma patient metagenomes **(Fig. 2a)**. A majority of patients with functional enterotype 2 were responders to anti-PD-1 therapy while most patients with functional enterotype 3 were non-responders **(Fig. 2b)**. Consistent with our other analyses, while functional enterotype 3 was characterized by a significantly higher abundance of total LPS biosynthesis genes **(Fig. 2c)**, functional enterotype 2 was characterized by a significantly higher abundance of *lpxM* **(Fig. 2d)**. Although direct measurements of lipid A acylation in complex mixtures like feces remains challenging, as a proof of principle, we utilized isolates from our culture collection^17^ and confirmed by LC-MS that gut commensal Gammaproteobacteria that encode *lpxM* can produce hexa-acylated lipid A **(Extended Data** Fig. 3).

**Figure 2.**
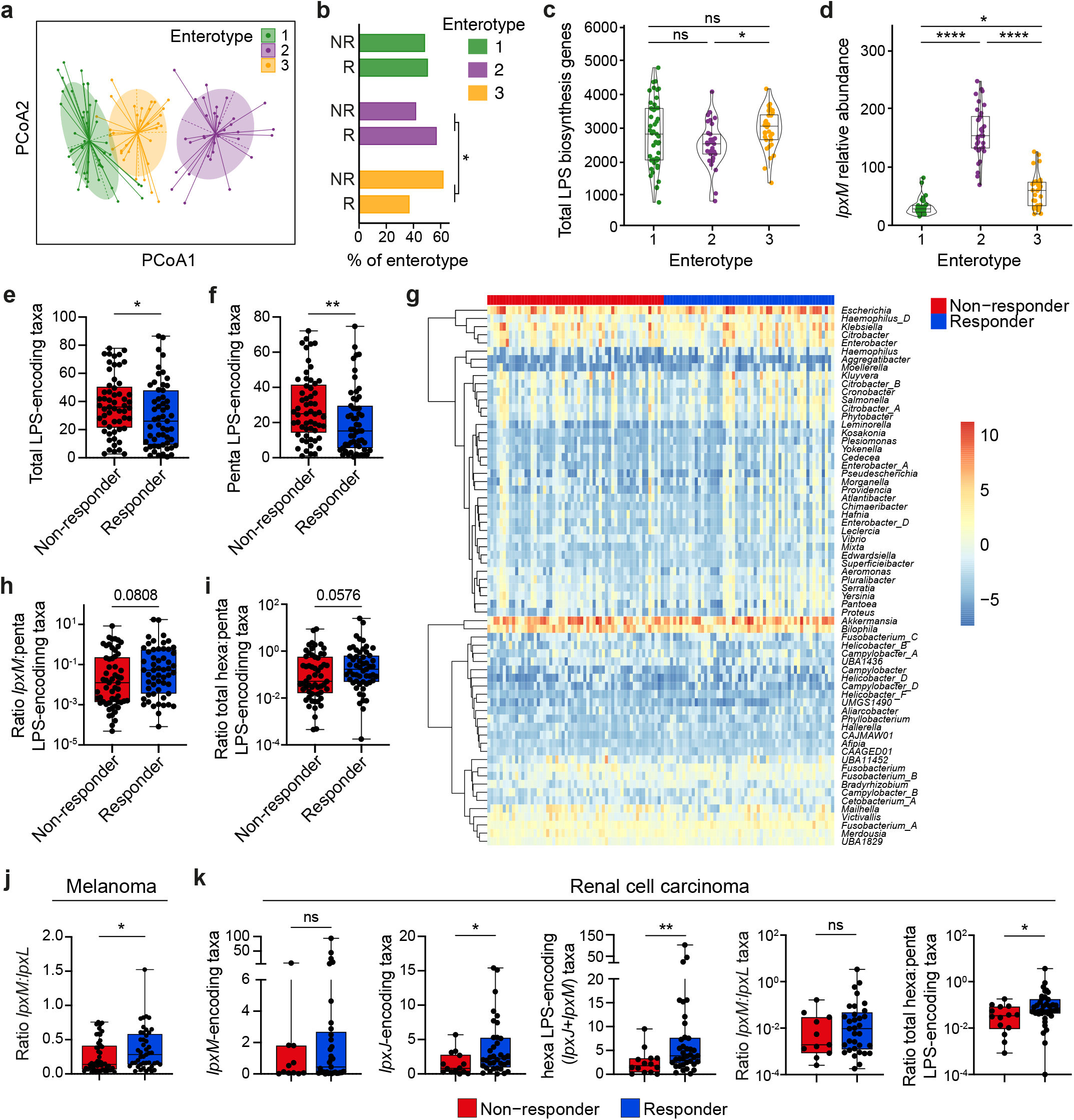
Functional metagenomic analyses indicate elevated hexa-acylated LPS ratios are favorable for patient responses to anti-PD-1 therapy. **a,** Functional enterotypes of metastatic melanoma patient gut microbiomes illustrated using Jensen–Shannon divergence (JSD) with partition around medoids clustering and principal coordinate analysis (PCoA). **b,** Percentage of non-responding (NR) and responding (R) patients within each enterotype. **c,** Relative abundances of total *lpx* encoding genes as count per million (CPM) and **d,** *lpxM* encoding genes within each patient enterotype. **e,** Total LPS-encoding taxa and **f,** penta-acylated LPS-encoding taxa as a percent of classified reads. **g,** Center-log ratio normalized relative abundance of *lpxJ-* and *lpxM*-encoding genera in patient fecal metagenomes clustered by rows (genera). **h,** Taxa predicted to produce *lpxM*- or **i,** *lpxJ-* and *lpxM*-mediated total hexa-acylated LPS as ratios to *lpxL*-encoding penta-acylated LPS taxa. **j,** Ratio of *lpxM* to *lpxL* genes based on CPM within fecal metagenomes of metastatic melanoma patients. **k,** Relative abundance as GCPM of (from left to right) *lpxM*, *lpxJ,* total hexa-acylation (*lpxM* and *lpxJ*) encoding genes and ratio to penta-acylation genes within fecal metagenomes of renal cell carcinoma patients prior to treatment with anti-PD-1. Metastatic melanoma patients, n = 112 (**a-j**). Renal cell carcinoma patients, n = 51 (**k**). One-tailed two proportions Z-tests without continuity correction (**b**). Kruskal-Wallis with post-hoc Dunn test and Bonferroni FDR correction (**c** and **d**). Mann-Whitney test with bars and error representing median and range (**e,f,h-k**). \**P* < 0.05, \*\**P* < 0.01, *\*\*\*\*P* < 0.0001.

Using taxonomic classification paired with pangenome annotation of LPS biosynthesis genes, we also quantified the abundance of microbial taxa that encode *lpxL* penta-acylated versus *lpxJ* and *lpxM* hexa-acylated LPS. On a taxonomic level, both total LPS-encoding and penta-acylated LPS-encoding taxa were significantly higher in non-responders to anti-PD-1 **(Fig. 2e,f)**. However, differential abundance of hexa-acylation was not dominantly mediated by a single genus or species of bacteria but rather a functional convergence on total LPS hexa-acylation capacity via multiple *lpxJ-* and *lpxM*-encoding taxa in the gut microbiota of responding patients **(Fig. 2g)**. This resulted in ratios of hexa- to penta-acylated LPS-producing taxa that were more than doubled in responders (mean *lpxM*-to-penta ratio: Responder = 0.9238, Non-responder = 0.4217 and mean total hexa-to-penta ratio: Responder = 1.255, Non-responder = 0.6244) **(Fig. 2h,i)** with significantly higher *lpxM*-to-*lpxL* ratios at a functional (gene or ORF) level **(Fig. 2j)**. Of note, we also functionally annotated and examined abundance of LPS biosynthesis genes in renal cell carcinoma (RCC) patients from a single previous study^2^. Although *lpxM* also trended higher in this smaller cohort, we found that in RCC patients *lpxJ*, which also mediates LPS hexa-acylation, was significantly more abundant in responders **(Fig. 2k)**, in line with the increased abundance of *Akkermansia muciniphila* previously noted in this patient cohort^2^. This also resulted in an increased ratio of total hexa-to-penta-acylated LPS genes in patients responding to anti-PD-1 **(Fig. 2k)**, raising the possibility that similar microbiome LPS signatures are associated with anti-PD-1 efficacy in other cancer types. Collectively, these data suggested that having an increased proportion of gut bacteria that encode hexa-acylated LPS is favorable for response to anti-PD-1 immunotherapy.

### Hexa-acylated LPS more potently stimulates host immunity

Given that the acylation state of LPS lipid A has been shown to directly impact immune signaling through TLR4^6,18^, we quantified the relative potency of hexa-acylated and penta-acylated LPS at stimulating NF-kB activation via TLR4 in a human monocyte/macrophage cell line reporter system. As expected, we found that TLR4 was required for LPS-mediated NF-kB activation while TLR2 was not, as deletion of TLR4, but not TLR2, significantly reduced NF- kB activity (**Fig. 3a**). Importantly, stimulation with hexa-acylated LPS resulted in substantially more activation than penta-acylated LPS (**Fig. 3a**). In contrast, expression of CD14, a receptor that also binds LPS but does not discriminate lipid A acylation states, did not affect differential responses to hexa- and penta-acylated LPS **(Extended Data** Fig. 4a). Hexa-acylated LPS purified from a Gammaproteobacterium was 10,000 times more potent at stimulating NF-kB activation than penta-acylated LPS from a Bacteroidota species **(Fig. 3b)**, resulting in higher cytokine secretion **(Fig. 3c).** This was blocked by the LPS-binding antibiotic polymyxin B (PMB) in a dose-dependent manner **(Extended Data** Fig. 4b). Notably, the addition of low, non-stimulatory concentrations of penta-acylated LPS significantly decreased the ability of hexa-acylated LPS to activate NF-kB **(Fig. 3d**), confirming that ratios of stimulatory to inhibitory LPS can dictate immune activation. To examine the effects of increasing the ratio of immunostimulatory hexa-acylated LPS in the gut, tumor-bearing mice were supplemented with a low dose of hexa-acylated LPS in their drinking water during anti-PD-1 treatment. Boosting levels of hexa-acylated LPS in the gut significantly improved the efficacy of anti-PD-1 therapy, resulting in decreased tumor burden **(Fig. 3e)** and significantly increased tumor infiltration of IFN-γ^+^ and TNF^+^ CD8^+^ T cells **(Fig. 3f)**. Collectively, these findings demonstrate that an increased relative abundance of hexa-acylated LPS in the gut may support the immune activation required for robust anti-PD-1 efficacy in cancer patients.

**Figure 3.**
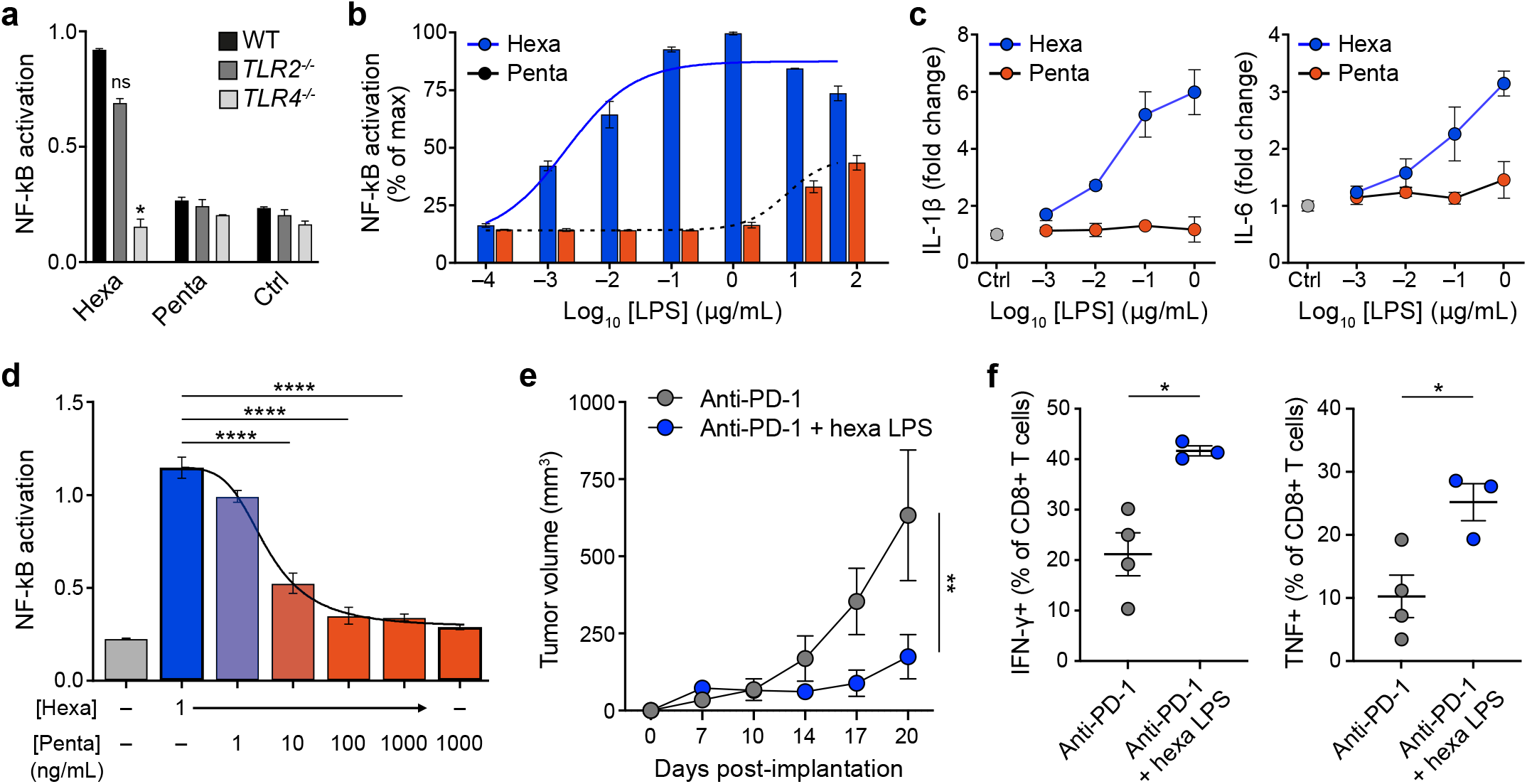
Increased ratios of hexa-acylated LPS more potently stimulate host immunity. **a-d,** Wild-type or *TLR4^-/-^* NF-kB reporter THP-1 (human monocyte/macrophage) cells were incubated with the indicated doses of hexa- or penta-acylated LPS for 24h. Data are representative of at least three experiments. **a,** NF-kB activation in response to 1ng/mL hexa- acylated LPS or 1 µg/mL of penta-acylated LPS. **b,** NF-kB activation dose-response with the indicated doses of hexa- or penta-acylated LPS. **c,** IL-1β (left) and IL-6 (right) secretion by THP-1 cells after 24h treatment with the indicated doses of LPS, shown as fold increase over media controls. **d,** NF-kB activation with the indicated combined doses of hexa- and penta- acylated LPS. **e,** MC38 colorectal adenocarcinoma cells were subcutaneously implanted into wild-type mice, and measurements were taken at indicated time points post implantation. Indicated animals were treated with anti-PD-1 i.p. and/or oral hexa-acylated LPS or control drinking water starting at day 10 after tumor implantation. Data shown are representative data from 1 of 2 experimental repeats with n = 4 mice per group. **f,** Flow cytometry measurements of frequency IFN-γ- and TNF-producing CD8^+^ T cells from tumors of animals in **e.** One-way ANOVA with a Dunnett’s multiple comparisons test (**d**), two-way ANOVA (**e**) and unpaired t test with Welch’s correction (**f**). Bars and error represent mean and s.e.m. \**P* < 0.05, \*\**P* < 0.01, *\*\*\*\*P* < 0.0001.

## Discussion

Improving response rates to ICI is essential for improving survival of patients. In this initial study, we functionally annotated a large cohort of patient microbiota profiles to uncover the role of gut bacteria in ICI efficacy. Although differences in patient population based on geographic location significantly contribute to variance at a taxonomic level (even after batch correction), this source of variation contributed far less at a functional level **(Extended Data** Fig. 1b,c), supporting the use of our functional metagenomics approach to see past taxonomic heterogeneity to an underlying mechanism involving LPS acylation. By exploring this mechanism *in vitro* and *in vivo*, we were able to demonstrate that hexa-acylated LPS increases immune activation in a TLR4-dependent manner, cautioning against use of polymyxin B and related antibiotics in immunotherapy patients and highlighting actionable potential avenues to therapeutically predict and pharmacologically improve patient outcomes.

It is important to note that inferred LPS biosynthesis capacity based on taxonomic identification and pangenome annotations did not always match our taxonomy-independent functional analyses at the level of LPS biosynthesis genes. This highlights potential limitations of taxonomy-based analyses for inferring function and may explain how meta-analyses in previous studies did not identify differential contributions of LPS hexa- and penta-acylation to immunotherapy responses. In addition, current databases and methods of functional annotation of microbiome genes are far from complete and comprehensive, necessitating our use of multiple approaches for both taxonomic and functional analyses in order to derive biological meaning.

Other microbiota-derived metabolites, including *Bifidobacterium* inosine^19^, *Enterococcus* muropeptides^20^ and bacterial flagellin^21^, have also been implicated in modulating ICI treatment response. While it is currently unknown how gut LPS signaling interfaces with signals from other microbiome-derived immunostimulatory metabolites, simultaneous activation of different pattern recognition receptors (PRRs) has been shown to result in synergistic immunostimulation^22,23^. It is therefore likely that multiple microbiota-mediated mechanisms that modulate treatment response co-exist and interact to define patient outcomes, perhaps contributing to the divergent taxonomic associations described by previous studies. In addition, host factors are also likely to play a role in this mechanism of immunotherapy modulation. For instance, differential expression of host intestinal alkaline phosphatase has been shown to affect the strength of the TLR4 signal^24^, and this represents just one of many currently uncharacterized patient factors that could be explored in the future.

Numerous other aspects of this microbiome-immunotherapy interaction also warrant further studies, including the contribution of TLR4 in different cell types to anti-PD-1 efficacy. In addition, methods to accurately measure different lipid A acylation states in complex environments like patient stool samples merit further development and optimization to determine the degree to which metagenomic profiles can reflect the heterogenous mixture of lipid A variants in the gut. A better understanding of the microbial metabolites and host signaling pathways that determine clinical response to ICI immunotherapies will enable design of new therapeutic approaches to overcome microbiota-driven heterogeneity in immunotherapy outcomes and better prognostication for the benefit of cancer patients.

## Methods

### Curation of publicly available datasets

We retrieved human metagenomic data from six publicly available datasets^1–4,16^ through the NCBI Sequence Read Archive using the accession numbers ERP127050, SRP339782, SRP197281, SRP116709, SRP115355, and ERP104577. These publicly available cohorts are shown in **Supplementary Table 1 and 2**. We included only those samples in our study which were collected at baseline i.e., before starting the therapy. We excluded any samples from patients who received combination therapy (i.e. anti-CTLA4 along with anti-PD1 or IFN*-*γ with anti-PD1) and/or had a recent history of proton pump inhibitor, H2 receptor antagonist or antibiotic usage whenever metadata for these parameters was available. We classified patients into responder and non-responder groups according to the outcomes reported in the respective studies above. Patients classified as stable disease (SD), partial response (PR) or complete response (CR) were considered responders, and patients with progressive disease PD were considered non-responders for the purposes of our analyses.

We are aware that working with public datasets may introduce biases and artificial variability into the analyses due to nonuniformity of the data structure. It is noteworthy that in our study we did not directly use the processed count or abundance matrices, which are known to contribute to batch effects. Along with the baseline filtering criteria mentioned above, we also implemented a uniform stringent quality control and downstream analysis method for all the samples at individual raw sequencing read levels, as detailed below. Thus, all the pooled sequencing samples were treated as part of one experiment, and the source of the sample (i.e. study and country) was used as a covariate, similar to the original studies.

### Quality control, and preprocessing

Pre-processing pipeline consists of three main stages: (1) initial quality control by removing low-quality reads (quality score <Q30), reads <75 bp and Illumina adapters from the forward and reverse reads using Trimmomatic software (v0.39)^1^. (2) DNA contamination removal using Bowtie 2 (v2.3.5.1)^2^ in sensitive mode, removing host DNA from humans (GRCh38) and non-host phiX174 Illumina spike-in. (3) Checking the FastQC (http://www.bioinformatics.babraham.ac.uk/projects/fastqc/) results and removing any samples that had >50% read duplication from the human-associated metagenomes.

### Contig assembly, gene-calling, and functional annotation

Quality-trimmed host decontaminated paired-end (PE) metagenomic reads were assembled individually using metaSPAdes (v3.15.4)3 to build contigs. Single-end (SE) samples (RCC cohort; **Supplementary Table 2**) were assembled using MEGAHIT (v1.2.9)^3^. Contigs from individual samples were concatenated into a single file and contigs smaller than 500 bp were discarded. To make a non-redundant contig database, 100% identical contigs were clustered and a representative longer contig was kept using cd-hit-est from CD-HIT program^4,5^. Prodigal (v2.6.3)^6^ was used in ‘meta’ mode for the open reading frame (ORF) and translated protein prediction from the non-redundant contig collection. The script sqm_annot.pl from SqueezeMeta software (v1.5.2)^7^ was used to assign KEGG orthology and corresponding microbial taxa to the predicted protein sequences. The latest databases were downloaded, using the script make_databases.pl from SqueezeMeta. In brief, functions were assigned using Diamond (v2.0.15.153)^8^ blastp alignments of the proteins against the KEGG^9^ and NCBI non-redundant (nr)^10^ databases using the lowest common ancestor^11^ methods. This way each protein sequence from the metagenomes was assigned to a function (KEGG) and bacterial taxa and thus a connection between function to taxonomy was established. To quantify the predicted genes in the samples across the metagenomes, we used the Salmon tool (v1.8.0)^13^. First, the predicted ORFs were indexed, and then reads (PE or SE) from individual samples were mapped onto the indexed Salmon database.

### Genomic prediction of bacterial LPS structure

To predict which bacterial species of the human gut microbiota produce LPS, *lpx* operon genes were identified from functional pangenomes of 3,006 bacterial species using the ‘feature_search’ module of the MGBC-Toolkit v1.0^17^. *lpx* genes were identified using both the InterPro and EggNOG functional schemata. Accessions for searched *lpx* genes are as follows:

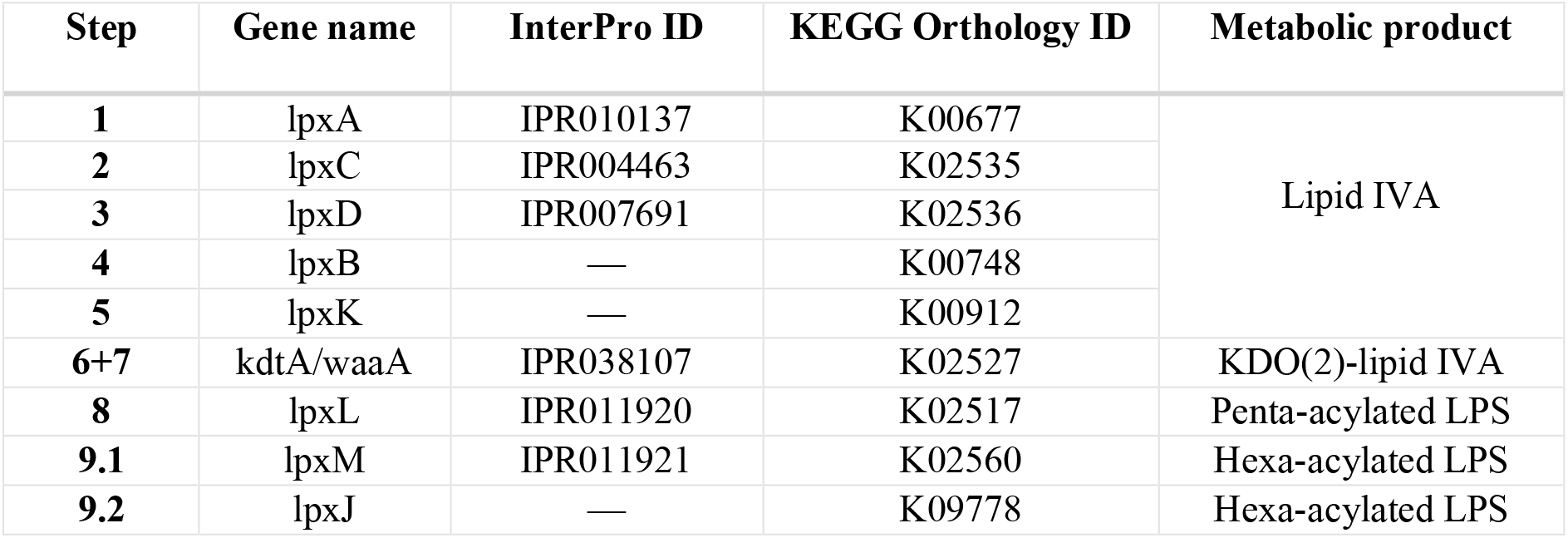

| Step | Gene name | InterPro ID | KEGG Orthology ID | Metabolic product |
| --- | --- | --- | --- | --- |
| 1 | *lpxA* | IPR010137 | K00677 | Lipid IVA |
| 2 | *lpxC* | IPR004463 | K02535 |  |
| 3 | *lpxD* | IPR007691 | K02536 |  |
| 4 | *lpxB* | — | K00748 |  |
| 5 | *lpxK* | — | K00912 |  |
| 6+7 | *kdtA/waaA* | IPR038107 | K02527 | KDO(2)-lipid IVA |
| 8 | *lpxL* | IPR011920 | K02517 | Penta-acylated LPS |
| 9.1 | *lpxM* | IPR011921 | K02560 | Hexa-acylated LPS |
| 9.2 | *lpxJ* | — | K09778 | Hexa-acylated LPS |

For each pangenome, genes that were encoded by at least 50% of the constituent genomes were considered to be encoded by that bacterial species. As most genes could be annotated using both InterPro and eggNOG, species that were identified as encoding a gene by at least one schema were included in analyses. All annotations were manually reviewed in context of the published literature and known structures for LPS. Following this review, only the InterPro database was used for *lpxM* annotation as eggNOG over-identified these genes in taxa which are known not to encode them.

LPS structure was then predicted using these annotation data according to the KEGG reference pathway for LPS biosynthesis. Species encoding *lpxACDBK* were considered to produce Lipid IVA and were thus defined as total Gram-negative LPS-encoding taxa (n=884) while taxa that additionally encoded *kdtA/waaA* were considered to produce KDO2-lipid IVA (n=827). Taxa that further encoded *lpxL* and either *lpxM* or *lpxJ* were considered to have the capacity to produce hexa-acylated LPS (*lpxLM*, n=127; *lpxLJ*, n=112), while taxa that further encoded *lpxL* but not *lpxM* or *lpxJ* were defined as penta-acylated LPS producers (n=538). LPS- encoding taxa that did not encode *lpxL* were defined as tetra-acylated LPS producers (n=107).

### Analyses of patient metagenomes

Metagenomes from melanoma and renal cell carcinoma patients were curated and quality-controlled as described above. For gene-level relative abundance, undetected (zero-value) samples were excluded and all non-zero values are reported. In renal cell carcinoma patients, the sample numbers for hexa-acylated LPS were as follows: *lpxM*, n=44; *lpxJ*, n=50, and total (*lpxM*+*lpxJ*), n=51. KEGG orthologous group (KEGG OG) present in less than 5 samples was considered as sporulating and or misannotation and removed from the analysis. Relative abundance of the predicted LPS biosynthetic genes (ORFs) from the patient metagenomes was compared between “Responder” and “Non-responder” through a single comparison and therefore there was no need for a multiple testing correction. Prior to functional analysis, we used ComBat-Seq ^25^ to correct the batch effects from the functional profile of the melanoma patient cohort. ComBat-Seq was run on the read count matrix with study metadata as batch and treatment response as group (full-model).

Taxonomic profiling was performed with Kraken v2.1.2 and Bracken v2.6.2 using a custom database built from representative genomes of the human gut microbiota20 that were re-annotated using the Genome Taxonomy Database v2.1. This database is available at doi.org/10.5281/zenodo.7319344. After analysis with Kraken2/Bracken2, we used ConQuR ^26^ to correct for batch effects between studies. ConQuR was run on the OTU read count matrix using a penalised fitting strategy (logistic_lasso=T, quantile_type="lasso", interplt=T) with study metadata as the batch variable and the response as a covariate. The resulting batch-corrected read count matrix was analysed in R.

### Data normalization

Gene abundance of the single melanoma^1^ and RCC cohort was expressed in the form of GCPM (Gene count per million).

The GCPM within the sample was calculated as: 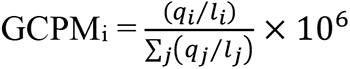 where qi denotes the number of raw reads mapped to the predicted gene, li is the gene length, and ∑_𝑗_(𝑞_𝑗_/𝑙_𝑗_) corresponds to the sum of mapped reads to the predicted gene normalized by gene length. Thus, GCPM accounts for gene length and sequencing depth and facilitates comparisons across samples. Since the batch correction was performed on the raw read counts of the pooled melanoma patient cohort, we used a slightly different method to normalize and scale our data. The batch-corrected count matrix was log-transformed and added with a pseudo count of 10^-6^ followed by scaling to count per million (CPM) value.

The CPM within the sample was calculated as: 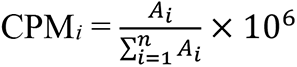 where *Ai* is the batch-corrected number of reads mapped to the predicted gene and *n* is the total number of predicted gene.

### Clustering and functional enterotype analysis

Functional enterotyping and between-class analyses were performed according to the procedure described by Arumugam et al.^15^. Data generated by functional annotation with the KEGG database (KEGG orthologs) were used to calculate the Jensen–Shannon divergence (JSD) among samples. The partitioning around medoids (PAM) clustering algorithm was applied to cluster the relative abundance of functional profiles of Genus or Orthologous groups (OGs). The number of optimal clusters was estimated using the Calinski-Harabasz (CH) index using the clusterSim package in R according to a previously described method^16^. The between- class analysis was performed to identify the major drivers of the metagenome functions and support clustering the OGs abundance profiles into clusters. The between-class analysis is a type of ordination with instrumental variables with the highest effect sizes that maximize the separation between classes of this variable. In this study, the instrumental variable was cluster classification using PAM clustering, obtained from KEGG OGs, which contributed the maximum to the principal coordinates, identified based on their eigenvalues. The analysis was performed using ade4 package in R.

### Mass spectrometry of acylation of lipid A from commensal gut bacteria

Mass spectrometry analyses of lipid A were performed using an Agilent 1290 Infinity II LC system coupled to an Agilent 6546 Q-TOF instrument. Lipid A-1P standard (Sigma Cat. No. L6638-1MG) was prepared in chloroform:methanol:water (74:23:3) at 1 mg/mL. 5 µL of 0.5 mg/mL of lipid A-1P was injected onto a Waters XBridge C18 column (50 mm x 2.1 mm, 3.5 µm particle size), and the mobile phase was a linear gradient of 0-95% isopropyl alcohol with 200 mM ammonium hydroxide and 5 µM medronic acid (mobile phase B) in methanol:water (80:20) with 200 mM ammonium hydroxide and 5 µM medronic acid (mobile phase A) over 17 min (before holding at 100% mobile phase A for 3 min). Flow rate was 0.25 mL/min and the column was operated at 45°C. Post-column addition of 10% DMSO in acetone at a rate of 0.25 mL/min was used. The published EICs by Sándor et al., 2016 ^27^ were used to characterize the lipid A molecule. Increases in acyl chain length led to later chromatographic elution.

Mouse commensal *E. coli* (Isolate: A7_7.EC.CDM; NCBI assembly GCA_910574425.1) and *K. pneumoniae* (Isolate: A5_5.KP.CDM; NCBI assembly GCA_910574035.1) from our mouse culture collection were confirmed to encode *lpxM* using the MGBC ToolKit with “feature_search” function. These isolates were inoculated in 5 mL of YCFA broth using a single colony from streaked YCFA agar plates. The cultures were incubated anaerobically overnight at 37 °C. The 5 mL cultures were used to inoculate 1L of YCFA liquid media. 1L liquid cultures were incubated anaerobically overnight at 37 °C to an OD600 of approximately 1.000. Cultures were separated into 200-mL tubes and centrifuged at 3,500xg for 20 minutes.

Media was discarded and bacterial cell pellets were washed once with PBS. Dry pellets were stored at -80 °C. The culture pellet was prepared at 100 mg/mL in water, hydrolyzed with a mild acid hydrolysis buffer (50 mM sodium acetate), and Bligh-Dyer extracted (proportion of chloroform:methanol:water, 1:2:0.8). Final resuspension was in chloroform:methanol:water (74:23:3) and 5 µL were injected onto the Waters XBridge C18 column (50 mm x 2.1 mm, 3.5 µm particle size), using the same chromatographic conditions as described above. The acyl chain lengths for the monophosphorylated and diphosphorylated forms of lipid A were annotated based on Sándor et al., 2016, and the proportional composition of these acyl chains was calculated for each isolate.

### In vitro cytokine secretion in monocyte/macrophages

THP-1 cells were cultured as indicated above, and supernatants were collected at 24h. RAW 264.7 cells were cultured in high Glucose DMEM (Sigma-Aldrich) with 10% fetal bovine serum (FBS, Sigma-Aldrich) and 100 U/mL Penicillin-Streptomycin (Gibco). 10^5^ cells per well were cultured for 24 hours in a final volume of 200uL culture medium with indicated concentrations of Ultrapure LPS from *E. coli* (hexa LPS, InvivoGen) and/or Ultrapure LPS from *P. gingivalis* (penta LPS, InvivoGen) with or without polymyxin-B (PMB, APExBio) in CELLSTAR 96 well plates (Greiner). Culture supernatants were mixed with the indicated cytokine beads and PE-conjugated detection antibodies from the Mouse Th1/Th2/Th17 CBA Kit (BD Bioscience) according to the manufacturer’s instructions. For THP-1 cells, the indicated cytokine cytometric beads and PE-conjugated detection antibodies from the BD Cytometric Bead Array (CBA) Human Inflammatory Cytokines Kit (BD Bioscience) were mixed with LPS treated THP-1 supernatants according to the manufacturer’s instructions. All samples were analyzed on a BD LSRFortessa™ Cell Analyzer (BD Bioscience).

### Mice

Nine-week-old wild-type C57BL/6 mice were used to perform *in vivo* tumor experiments. Mice were purchased from Charles River and housed at the University Biomedical Services (UBS) Anne McLaren Building (Cambridge). Experiments were conducted in accordance with UK Home Office guidelines and approved by the University of Cambridge Animal Welfare and Ethical Review Board.

### Tumor challenge and treatment

MC38 colon carcinoma cells were purchased from Kerafast. Cell lines were passaged in DMEM (Sigma-Aldrich, D5796) supplemented with 10% FBS (Sigma, BCCC3714), 1% Glutamax (ThermoFisher, 35050-038), 1% Non-essential amino acids (NEAA) (ThermoFisher, 11140-035), 1% Sodium pyruvate (ThermoFisher, 11360-039), 1% Penicillin-Streptomycin (ThermoFisher, 15140-122), and 0.1% of 2-mercaptoethanol (ThermoFisher, 21985023), Amphotericin B (ThermoFisher, 15290-026), Gentamycin (ThermoFisher, 15750- 045). Mice under isoflurane anaesthesia were injected subcutaneously (s.c.) with 10^6^ cells in 100 µl of sterile PBS into the right flank. Tumors were measured using a caliper at day 7, 10, 14, 17, and 20 after implantation. Mice were injected intraperitoneally (i.p.) with 200 µg of anti-PD-1 (clone 29F.1A12) (BioXcell, BE0273) or IgG2a isotype control (BioXcell, BE0089) twice per week. Mice were administered 25mg/L hexa-acylated LPS (Sigma, L2630-100MG) in sterile drinking water 10 days after tumor implantation. LPS-containing water was changed twice per week.

### Sample processing of murine tumors for immunophenotyping

Tumor samples were digested in RPMI containing collagenase D and DNase1 at 37°C for 30 minutes. A 40/80 Percoll gradient was used to isolate lymphocytes from tumors. Cell suspensions were filtered using 40-µm cell strainers.

### Flow cytometry of tumor-infiltrating lymphocytes and cytokines

Isolated lymphocytes were re-stimulated with eBioscience™ Cell Stimulation Cocktail (plus protein transport inhibitors) (500X) (Invitrogen, 00-4975-93) in complete RPMI with 10% FBS in a 37°C cell culture incubator with 5% CO2 for 4.5 hours. Dead cells were then stained using the LIVE/DEAD™ Fixable Aqua Dead Cell Stain Kit (Thermo Fisher Scientific, L34965). Cells were stained with surface/extracellular antibodies (Table 1) for 20 minutes on ice, and permeabilized for 30 minutes using the BD Cytofix/Cytoperm™ Fixation/Permeabilization Kit (BD Biosciences, 554714) according to the manufacturer’s protocol. Intracellular antibodies (Table 1) were then added for 30 minutes. A Cytek® Aurora flow cytometer was used to analyze samples. Data were analyzed using FlowJo™ Software (v10.8.1).

**Table 1.** Re-stimulation Flow Cytometry Panel

| Antibody | Fluorescence | Clone | Catalog No. |
| --- | --- | --- | --- |
| Extracellular Antibodies |  |  |  |
| CD45 | BUV805 | 30-F11 | Thermo, 368-0451-80 |
| TCRβ | APC-eFluor 780 | H57-597 | Thermo, 47-5961-82 |
| TCR$\gamma\delta$ | FITC | GL-3 | Thermo, 11-5711-82 |
| CD4 | PE-Cy7 | RM4-5 | Thermo, 25-0042-82 |
| CD8α | BUV737 | 53-6.7 | Thermo, 367-0081-82 |
| Intracellular Antibodies |  |  |  |
| IFNγ | Percp-Cy5.5 | XMG1.2 | Thermo, 45-7311-82 |
| TNFα | APC | MP6-XT22 | Thermo, 17-7321-82 |

### Statistical Analysis

Statistical analyses were performed using either Graphpad Prism software or R. Non-parametric Kruskal-Wallis tests were used as indicated to calculate statistical significance of the difference in sample means and post-hoc Dunn *p* values with Bonferroni FDR adjustment were calculated for the pairwise comparisons among the enterotypes. The proportions of clinical responders were compared between enterotypes using one-tailed two proportions Z-tests without continuity correction. Two-way ANOVA was used to evaluate statistical significance in experiments with tumor growth measurements. We used Non-metric Multi- Dimensional Scaling (NMDS), a distance-based ordination technique [https://doi.org/10.2307/1939814, https://doi.org/10.1111/j.1442-9993.1993.tb00438.x] in R (`vegan::metaMDS()`) with default `engine = "monoMDS"`. In the process of calculation of goodness-of-fit, a single NMDS run identifies a local minimum therefore we iterated the runs 1000 times, or the best solution achieved to reach the global minimum (convergence) using the function Procrustes in R (`vegan:: procrustes()`). The final stress score was used to assess the optimal model and goodness-of-fit as suggested previously [https://doi.org/10.1007/BF02289565, https://doi.org/10.1007/BF02289694]. In addition to the stress score, we also performed a two-tailed *t-test* on the NMDS coordinates to highlight the difference between the “Responder” and “Non-responder” groups. *P* values of less than 0.05 were considered statistically significant. Statistical tests used are specified in the figure legends. *P* values correlate with symbols as follows: ns = not significant, * *P* ≤ 0.05, ** *P* ≤ 0.01, *** *P* ≤ 0.001, **** *P* ≤ 0.0001.

## Acknowledgements

V.A.P. is supported by a Sir Henry Dale Fellowship jointly funded by the Wellcome Trust and the Royal Society [206245/Z/17/Z]. B.S.B. was supported by a studentship from the Rosetrees Trust [A2194]. We thank members of the Pedicord, Cross, Bryant, Roychoudhuri and Okkenhaug laboratories for sharing of reagents, protocols, ideas and discussion.

