## Extended Data Fig for "Hexa-acylated lipopolysaccharides from the gut microbiota enhance cancer immunotherapy responses" (from another version of this preprint: version 1, 2024-06-25)

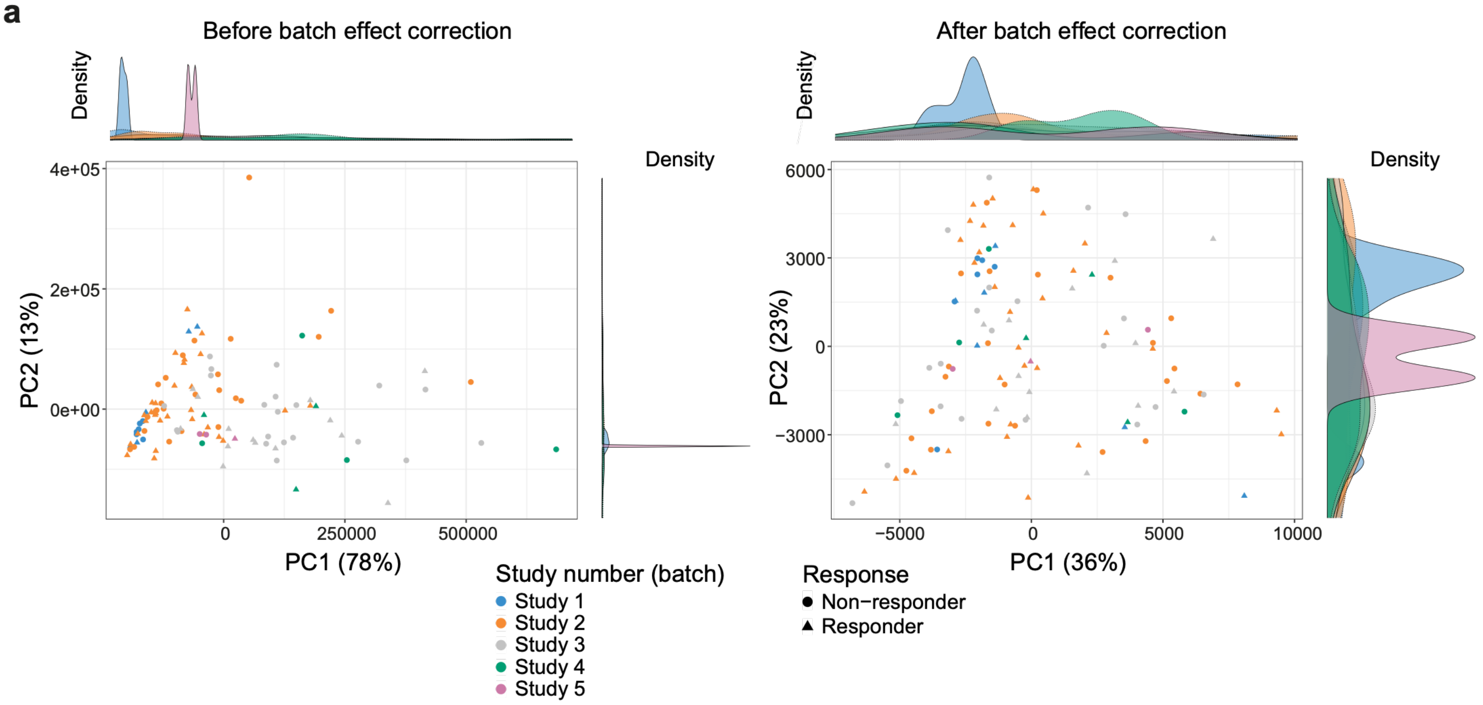


**b**

Function **before** batch correction (PERMANOVA):

| Source of Variation | Df | Sum Sq | $R^2$ | F | Pr(>F) |
|---|---|---|---|---|---|
| **Study** | 4 | 2.9381 | 0.24504 | 9.0969 | 0.001 *** |
| **Country** | 2 | 0.5535 | 0.04617 | 3.4277 | 0.005 ** |
| **Response** | 1 | 0.1011 | 0.00843 | 1.2519 | 0.248 |
| **Residual** | 104 | 8.3974 | 0.70036 | | |
| **TOTAL** | 111 | 11.9901 | 1.00000 | | |

Function **after** batch correction (PERMANOVA):

| Source of Variation | Df | Sum Sq | $R^2$ | F | Pr(>F) |
|---|---|---|---|---|---|
| **Study** | 4 | 0.01402 | 0.00522 | 0.1429 | 1.000 |
| **Country** | 2 | 0.09764 | 0.03635 | 1.9903 | 0.060 |
| **Response** | 1 | 0.02339 | 0.00871 | 0.9537 | 0.351 |
| **Residual** | 104 | 2.55097 | 0.94972 | | |
| **TOTAL** | 111 | 2.68602 | 1.00000 | | |

**c**

Taxonomy **before** batch correction (PERMANOVA): :

| Source of Variation | Df | Sum Sq | $R^2$ | F | Pr(>F) |
|---|---|---|---|---|---|
| **Study** | 3 | 27686 | 0.03158 | 1.4724 | 0.031 * |
| **Country** | 3 | 188245 | 0.21470 | 10.0113 | 0.001 *** |
| **Response** | 1 | 9011 | 0.01028 | 1.4376 | 0.105 |
| **Residual** | 104 | 651850 | 0.74345 | | |
| **TOTAL** | 111 | 876792 | 1.00000 | | |

Taxonomy **after** batch correction. (PERMANOVA):

| Source of Variation | Df | Sum Sq | $R^2$ | F | Pr(>F) |
|---|---|---|---|---|---|
| **Study** | 3 | 9242 | 0.00987 | 0.3977 | 1.000 |
| **Country** | 3 | 108719 | 0.11609 | 4.6788 | 0.001 *** |
| **Response** | 1 | 12979 | 0.01386 | 1.6757 | 0.039 * |
| **Residual** | 104 | 805539 | 0.86018 | | |
| **TOTAL** | 111 | 936479 | 1.00000 | | |

**Extended Data Figure 1. Batch corrections reduce variance from study and geographic location to reveal microbiome associations with anti-PD-1 response. a,** Principal component analysis of the functional profile (KEGG OGs) of the pooled melanoma cohort before and after batch correction. Individual studies are shown in different colors. Histograms with different colors on the axes represent the sample distribution per study. Before batch correction samples were mostly clustered by study (left) while batch correction removed that effect (right). **b,** Permutational multivariate analysis of variance (PERMANOVA) results from the functional annotation before and after the batch correction. On the functional level, studies collectively accounted for 24.5% of the total variance before batch correction (p=0.001), while after batch correction, only 0.52% of the variance was attributed to the study. **c,** PERMANOVA results from the taxonomic annotation before and after the batch correction. *$P < 0.05$, **$P < 0.01$, ***$P < 0.001$

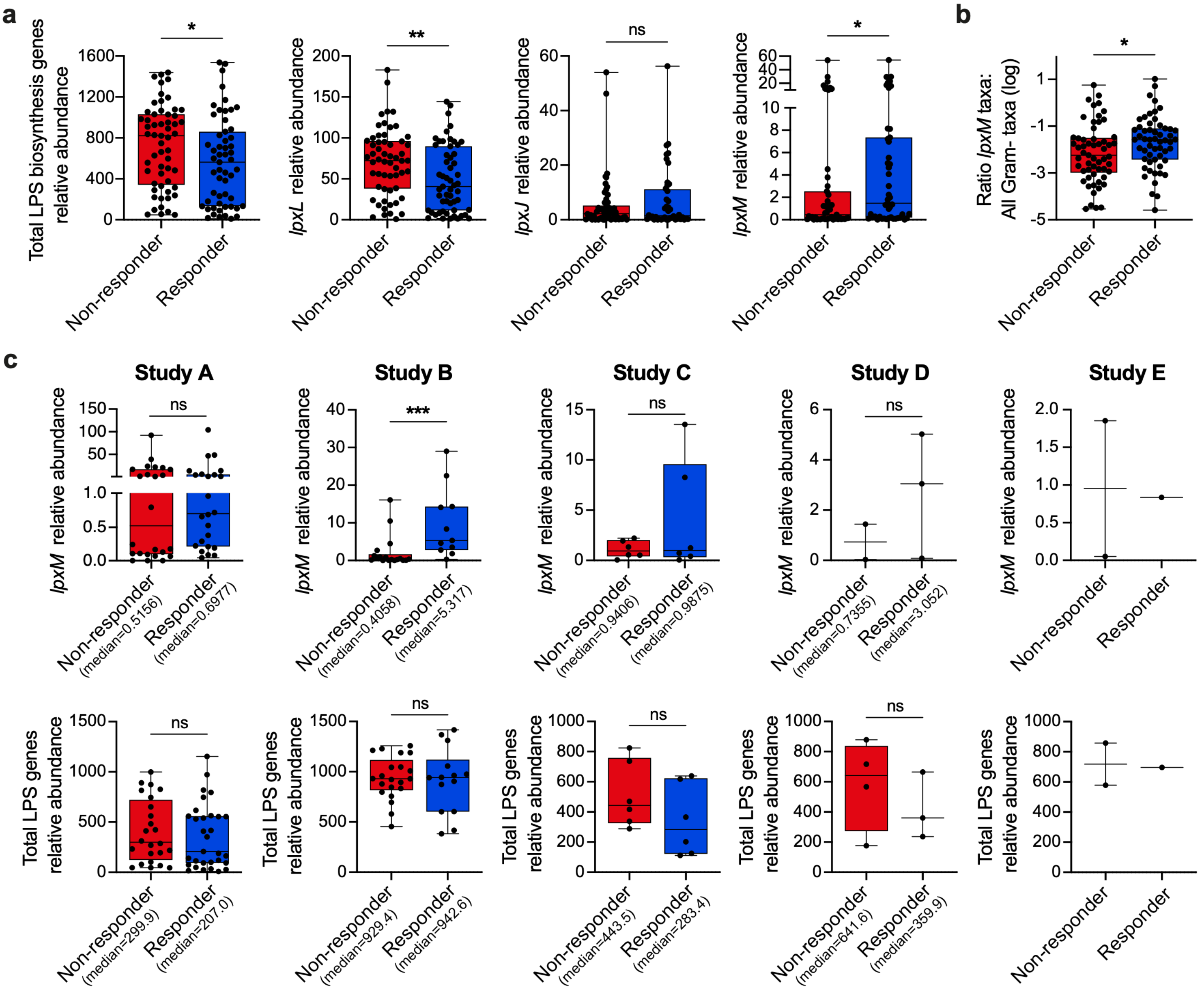


**Extended Data Figure 2. Relative abundances without batch correction of LPS-encoding genes in melanoma patient metagenomes prior to anti-PD-1 treatment. a,** Relative abundance as gene count per million (GCPM) of total LPS biosynthesis genes, *lpxL*, *lpxJ*, and *lpxM*, respectively (from left to right) in responder and non-responder metastatic melanoma patients. **b,** Ratio of *lpxM* encoding taxa to all other gram-negative taxa in indicated patient metagenomes. **c,** Relative abundance as GCPM of *lpxM* (upper row) and total LPS biosynthesis genes (lower row) within metagenomes of responder and non-responder metastatic melanoma patients in individual studies. Mann-Whitney test with bars and error representing median and range. $^*P < 0.05$, $^{**}P < 0.01$, $^{***}P < 0.001$.

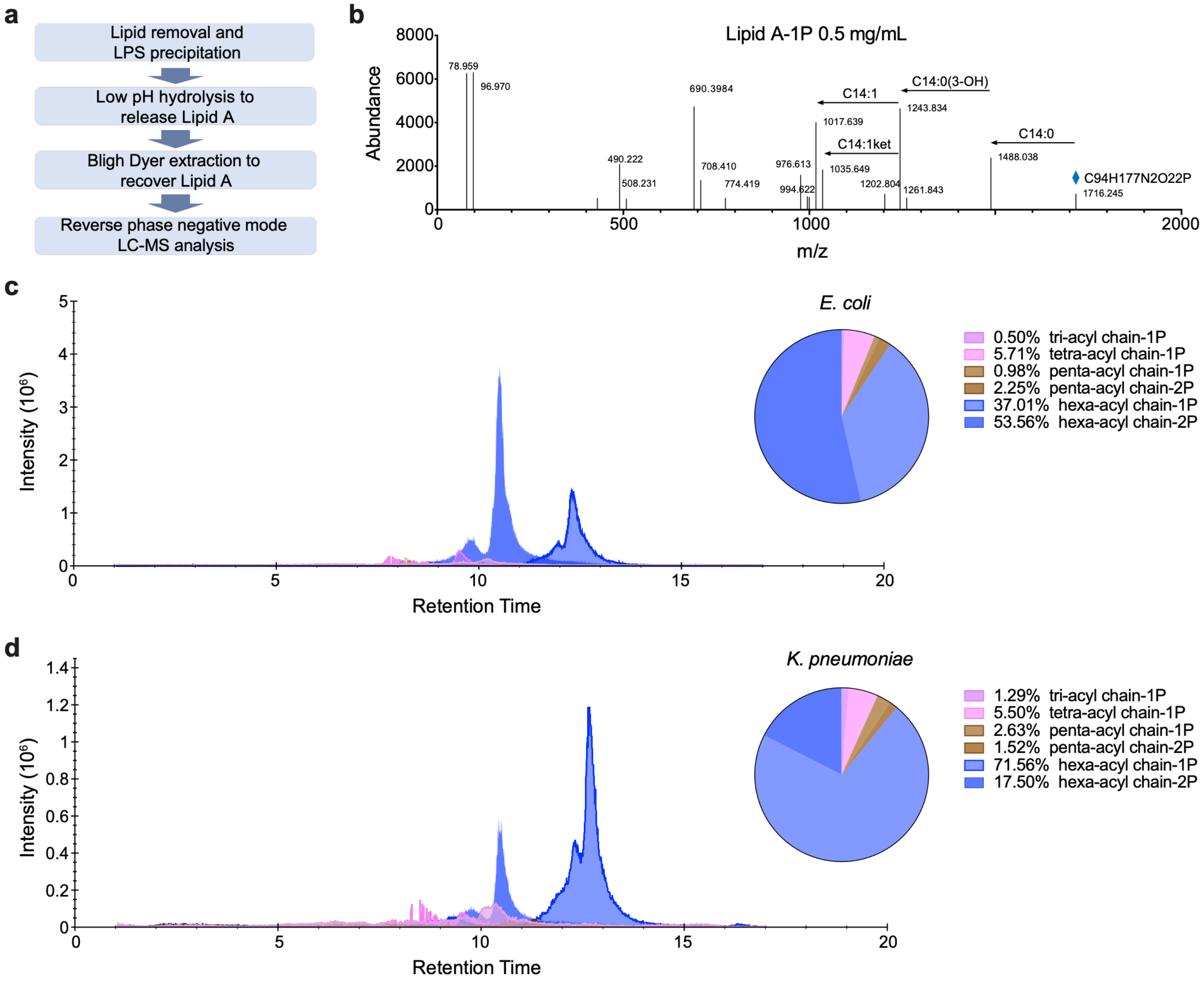


**Extended Data Figure 3. Genomically encoded *lpxM* accurately predicts lipid A hexa-acylation capacity of gut microbes.** Two commensal isolates from our culture collection that encode *lpxM* were grown anaerobically, and **a,** lipid A was isolated from resulting cell pellets and analyzed using a customized LC-MS lipidomics method for characterization of lipid A acylation. **b,** MS/MS spectra of one characteristic hexa-acyl-1P ion (*m/z* 1716) of a purchased standard lipid A-1P from *Escherichia coli*. The characteristic acyl chain losses are annotated in the spectra as described by Sándor et al., 2016. **c,** Overlaid extracted ion chromatogram (EIC) of lipid A from mouse *E. coli* isolate LPS (left) and relative composition in terms of acylation and phosphorylation (right). **d,** Overlaid EIC of lipid A from mouse *Klebsiella pneumoniae* isolate LPS (left) and relative composition in terms of acylation and phosphorylation (right).

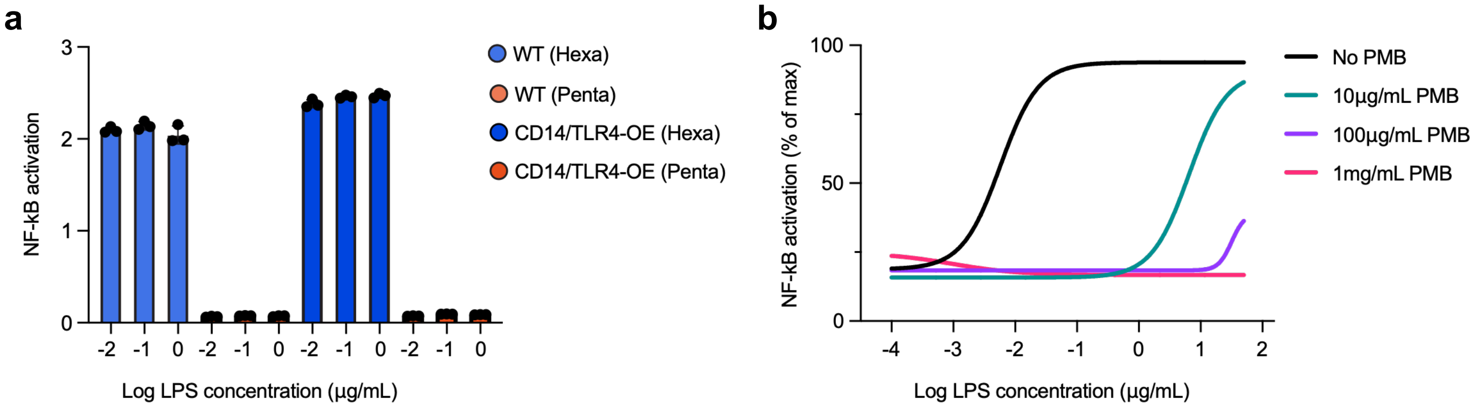


**Extended Data Figure 4. CD14 does not discriminate LPS acylation states, and polymyxin B blocks hexa-acylated LPS-mediated immune activation.** **a,** NF-kB reporter THP-1 cells that do not (WT) or do over-express CD14 and TLR4 (CD14/TLR4-OE) were incubated with the indicated doses of hexa- or penta-acylated LPS for 24h. Technical triplicates are plotted in representative plots from one of two experiments. No significant differences between WT and CD14/TLR4-OE were calculated by Mann-Whitney tests comparing equivalent doses and types of LPS. **b,** WT NF-kB reporter THP-1 cells were incubated with the indicated doses of hexa-acylated LPS and polymyxin B (PMB) for 24h. NF-kB activation is expressed as a percentage of maximum. The means of technical duplicates are plotted in representative plots from one of three experiments.
